# Genes Underlying Adaptive Physiological Shifts Among Hibernating Mammals

**DOI:** 10.1101/2024.10.15.618548

**Authors:** Danielle H. Drabeck, Myana Anderson, Emma Y. Roback, Elizabeth R. Lusczek, Andrew N. Tri, Jens Flensted Lassen, Amanda E Kowalczyk, Suzanne McGaugh, Tinen L Iles

**Affiliations:** University of Minnesota, College of Biological Sciences, Department of Ecology, Evolution, and Behavior; University of Minnesota, Medical School, Department of Surgery; Minnesota Department of Natural Resources, Forest Wildlife and Populations Research Group; Department of Cardiology B, Odense University Hospital Odense, Denmark; Form Bio, Austin, TX

## Abstract

Hibernation has evolved several times in mammals to overcome harsh winter climates and food scarcity. During hibernation, animals undergo extreme shifts in metabolic rate, heart rate, respiration, and body temperature. These changes reduce energy consumption and allow animals to survive solely on their fat reserves. Understanding the mechanisms for these extreme shifts has long been recognized as a model for translational medicine as hibernators do not exhibit the same adverse effects of extended immobility that non-hibernating mammals suffer. Though work on individual species has illuminated important mechanisms of these functional changes, the genomic basis of this phenotype remains largely unknown, and few studies have drawn on comparative work to elucidate commonalities across diverse hibernating mammals. Synthesizing both single species and comparative approaches, we use metabolomic data from active and denning black bears (*Ursus americanus*) to guide bioinformatic analyses of genes using tests of selection and evolutionary rate convergence across independent lineages of hibernating mammals. We identify several genes with significant signatures of selection and evolutionary rate convergence in hibernators that represent both previously known and novel genetic mechanisms of the hibernation phenotype. These data provide novel insights into the genetic basis of this adaptation and serve to direct clinical research in hibernation-based therapies.

## Introduction

Mammalian hibernation is a complex behavioral and physiological trait that has arisen in no fewer than 11 independent lineages of mammals and facilitates survival through extreme conditions (low temperature and food scarcity) during winter ^1,2^. Hibernation is characterized by a seasonal prolonged reduction in metabolic rate, body temperature, and heart rate, which rebounds to normal levels during waking months. During hibernation, animals exhibit considerable physiological shifts that allow them to survive low temperatures and go without food, water, or substantial movement for months^1^. Characterizing the physiological changes that occur during hibernation has been of central interest not only to evolutionary biology, but also to various applied medical fields (traumatic injury, long sedation on ventilators, coagulopathy, osteoporosis, muscular atrophy, and diabetes), and even space travel ^3–10^.

Metabolic parameters of hibernating mammals indicate suppression of mitochondrial respiration, upregulation of lipid oxidizing enzymes, and concomitant downregulation of glycolytic enzymes^11–13^. This metabolic reprogramming enables hibernators to use alternative fuels such as fatty acids and ketone bodies when glucose availability is low ^12^. Additionally, mammals have a complex regulation of hormones that influence their metabolism during hibernation. For example, iodothyronamine, leptin, and ghrelin may play a role in modulating thermogenesis, energy expenditure, and appetite ^4^. Previous work examining metabolic shifts in hibernators has led to important insights into the proteomic and genomic basis of basic physiological mechanisms that are widely applicable, for example, identification of heat shock protein 47 (HSP47) as a vital thrombotic regulator ^14^. Hibernation research has also revealed critical insights into mammalian physiology, offering a deeper understanding of metabolic suppression, resistance to tissue damage, and muscle preservation during prolonged inactivity ^15^. These discoveries have led to important medical applications, including therapeutic hypothermia for ischemic injury management and strategies to prevent muscle atrophy in immobilized patients ^16–20^. Besides therapeutic hypothermia, hibernation-based therapies have become an ever-expanding field as research has shown the efficacy of chemically inducing the same or similar pathways and physiologies seen during hibernation ^6,21,22^.

While metabolic studies have advanced our understanding of the physiological mechanisms underlying hibernation, the genetic basis of these adaptations—and whether they are conserved across hibernating species—remains largely unknown. This significant gap in knowledge limits our ability to manipulate these key physiological processes for potential therapeutic applications. Understanding mechanisms shared by all hibernating mammals not only sheds light on the repeatability of complex adaptations but also highlights evolutionary targets that are more likely to be conserved across all mammals, making them promising candidates for human medical applications.

Using plasma samples from American black bears (*Ursus americanus*), we find that a key metabolite (carnitine) shifts during the summer (active time), early denning (fully hibernating), and late denning (transitioning to waking) periods. We expand on this insight by using bioinformatic tools to build a dataset of genes related to carnitine. We also integrate a physiology-agnostic analysis of genes from 120 mammalian genomes, to identify genes evolving at convergent rates in hibernators. Together, our data reveal genes evolving under selection and at convergent evolutionary rates among more than 20 hibernating mammals that represent at least nine independent shifts to hibernation (Figure 1). By combining single-species metabolic insights with a comparative phylogenetic approach, our work identifies specific genes that may drive critical shifts in metabolite abundance in hibernating mammals. Our data elucidate novel genetic targets of selection in hibernating mammals, as well as confirm the importance of genes identified in prior studies of expression, proteomics, and predictive genomics. This work highlights the value of integrating both top-down and bottom-up methods and contributes vital insights into the genetic basis of physiologies that facilitate hibernation, providing promising avenues to translational medicine aiming to both understand and manipulate the same pathways and physiological processes.

**Figure 1.**
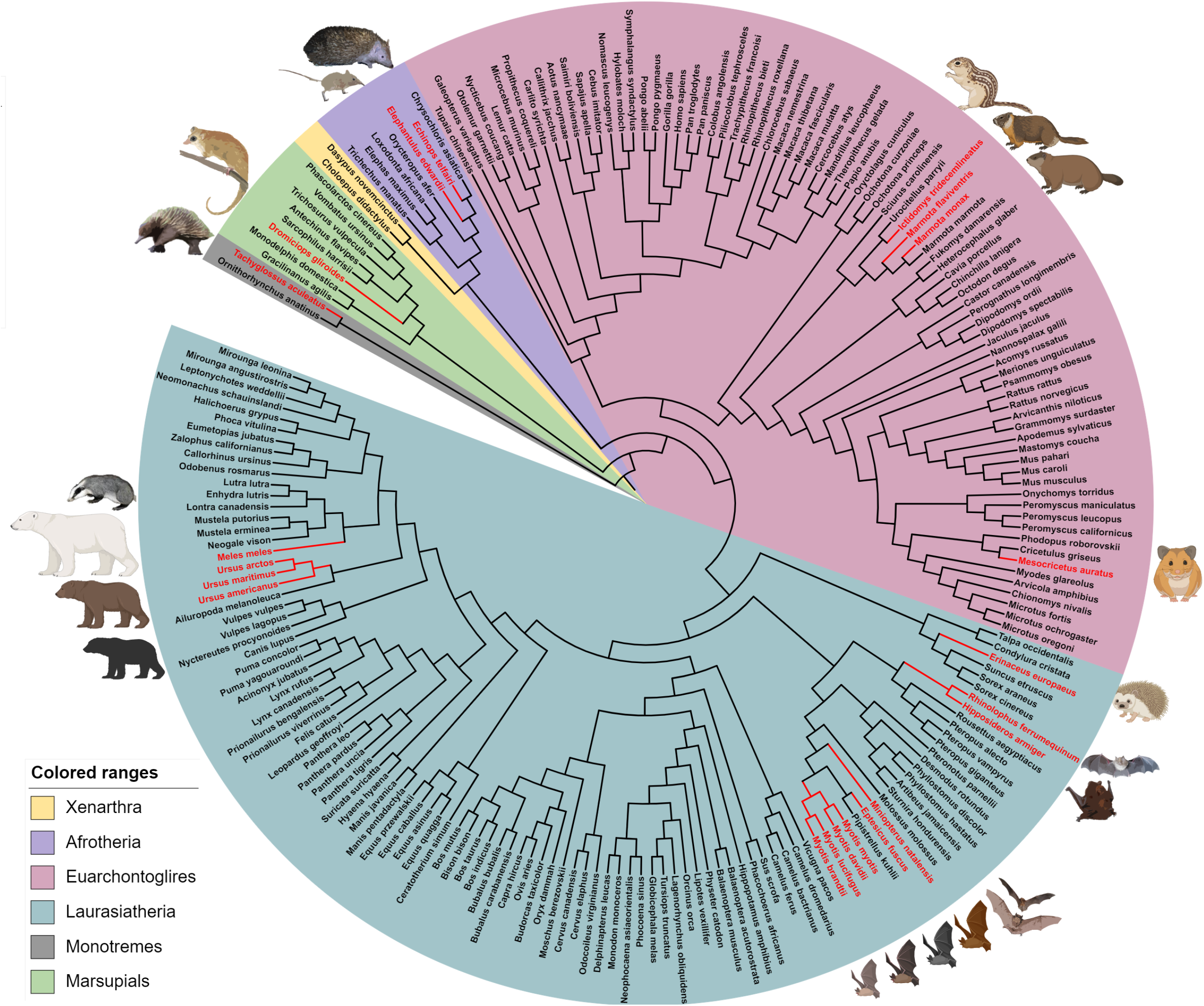
Rooted topology showing phylogenetic relationships of the mammals used for genomic comparative analyses. Lineages known to hibernate are indicated in red.

## Methods

### Bear Sample Acquisition

Free-ranging black bears are studied by the Minnesota Department of Natural Resources (MNDNR) to better understand the population dynamics and track the health and biometrics of the population. For this study, we collaborated with the MNDNR to collect blood samples to research black bear adaptations and physiology. Venous blood samples were collected while bears were anesthetized during den visits at three seasonal time points: 1) the active time during the summer months (n=12, June and July), 2) early-denning, their most immobile time (hibernations, November, December, January, n=35); and 3) late-denning as they are emerging from hibernation (February, March, n=28). Bears were anesthetized using standard wildlife handling procedures^23,24^. Biometric data were gathered, including height, weight, and the presence of cubs/yearlings (Supplementary Table 1) Whole blood was collected in citrated plasma tubes and spun for plasma and stored at -80C.

### Identifying Key Metabolic Indicators

All metabolites were identified using isotopically labeled internal standards and multiple reaction monitoring. Multiple reaction monitoring is a targeted mass spectrometry technique that selectively detects specific ion transitions for precise quantification. This method allows for the precise identification and quantification of metabolites under optimized conditions provided by Biocrates (Innsbruck, Austria), ensuring consistent and reliable results for metabolomic analysis. Isotopically labeled internal standards are added to ensure accurate quantification and account for sample handing and preparation differences. For quantification either a seven-point calibration curve or one point calibration was used depending on the metabolite class. Data were captured using the Analyst (Sciex) software suite and transferred to the MetID software (version Oxygen; Biocrates Life Sciences AG, Innsbruck, Austria) which was used for further data processing and analysis. Metabolites with less than 80% detection between all experimental samples were removed from our dataset, leaving 385 metabolites for our analysis. Any missing values were imputed using a K-nearest neighbor approach (k = 7) ^25^. The entirety of the dataset was normalized using quantum normalization methods in the preprocessCore R-package ^26^. The samples were categorized into three time periods: summer, early denning, and late denning. Data was subset by liquid chromatography methods and flow injection analysis for broad analysis of proteins.

Metabolic indicators are derived from formulas, such as ratios or sums of metabolite abundances, transforming raw metabolomic data into interpretable metrics that provide insights into biological processes. Raw metabolic abundance data was used to calculate 159 predefined metabolic indicators using the Biocrates MetaboINDICATOR™ software tool ^27,28^. To identify differences in metabolic indicators by denning period (linked to hibernation), we performed a sparse partial least squares discriminant analysis (sPLS-DA) using the MixOmics R package^29^. To evaluate sPLS-DA model performance and feature stability, we performed a three-fold cross-validation (CV) 10 times for each type of prediction distance (max distance, centroids distance, and Mahalanobis distance). Slight differences between the overall error rate and the balanced error rate (BER) (with BER being higher) were observed, indicating potential misclassification of minority classes. To assess the robustness of the model, we performed a feature stability analysis, leading to the selection of components 1 and 2 for the final PLS-DA model. This analysis was repeated with only early and late denning clusters to verify the metabolic indicators which showed the largest differences between these two phases (Figure 2A &B).

**Figure 2.**
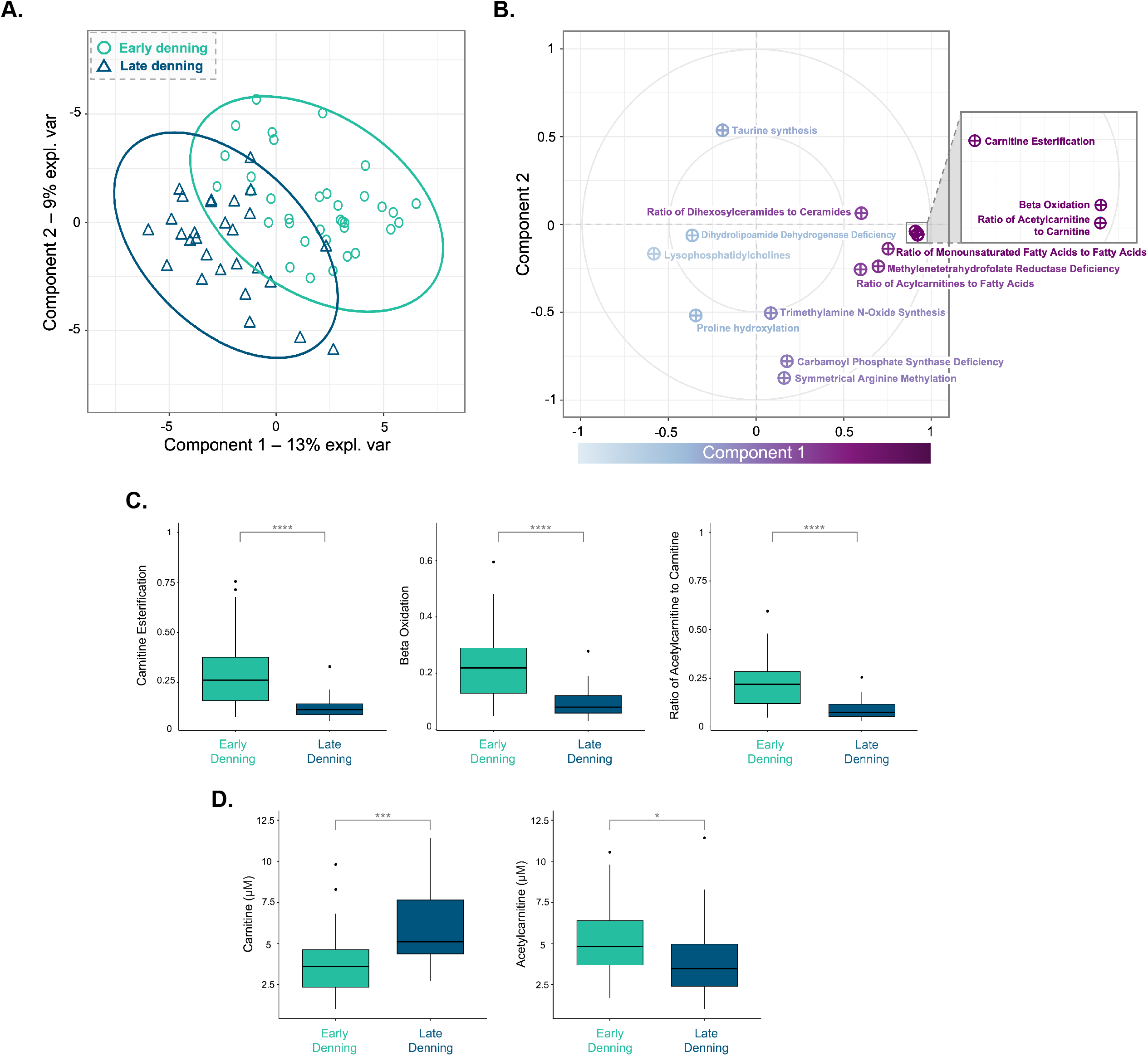
A. Sample plot generated using Sparse Partial Least Squares Discriminant Analysis (sPLS-DA) with the *mixOmics* package in R. The samples are colored according to their denning period and ellipses represent a 95% confidence interval. Components 1 and 2 explain 13% and 9% of the variation, respectively. B. Circle plot visualizing the correlation between metabolic indicators and the latent variables (Component 1, Component 2) derived from sPLS-DA. Each point represents a metabolic indicator, with the angle indicating its contribution to the direction of cluster separation and the distance from the center representing the strength of its correlation with the latent variables. Metabolites further from the center have a stronger influence on cluster separation, guiding the clusters in the directions shown. C. Metabolic indicators between early and late denning identified to have the strongest correlation with latent variables. All metabolic indicators are unitless quantities derived from Biocrates MetaboINDICATOR™: MxP® Quant 500 formulas. D. Raw metabolite abundance for Carnitine and Acetylcarnitine, the metabolites with <80% valid quantitation that contribute to the three metabolic indicators found using sPLS-DA.

### Identifying Key Metabolites

To identify metabolites underlying differences in metabolic indicators between early and late denning (hibernation vs waking), we further scrutinized the metabolic indicators identified in sPLS-DA analysis which appeared to have the starkest differences between late and early denning (Figure2B) (beta-oxidation, ratio of acylcarnitine to carnitine, and carnitine esterification). We confirmed these indicators were significantly different between denning periods using a Kruskal-Wallace test of significance followed by a Dunn’s pairwise test with Bonferroni multiple-comparison correction (p<0.05) (Figure 2C). For all three metabolic indicators, we used the provided MetaboINDICATOR™: MxP® Quant 500 formulas to identify raw metabolites used to derive the indicator variables. We used the same method (Kruskal-Wallace test of significance followed by a Dunn’s pairwise test with Bonferroni multiple-comparison correction) to find which metabolites were significantly different between early and late denning (p<0.05). Of these, only Acetylcarnitine (C2) and carnitine (C0) were above the 80% valid quantitation cut-off. When examining raw abundance only carnitine (C0) was identified as significantly different between early and late denning after multiple test correction (Figure 2D; Supplementary Figure 1).

### Genetic Data Curation

To identify a list of genes potentially underlying changes in carnitine levels during hibernation, genes related to pathways that involve carnitine were selected through the Kyoto Encyclopedia of Genes and Genomes ^30^. Through KEGG, carnitine metabolism was found to be present in the following pathways: thermogenesis, fatty acid degradation, insulin resistance, diabetic cardiomyopathy, and bile acid secretion. Using the ortholog table available for each pathway on KEGG, we searched NCBI’s ortholog database for multi-species ortholog sequence data. We identified 394 genes that were available for mammals, and the associated RefSeq transcripts and protein alignments were downloaded through NCBI’s ortholog database.

### Alignment and Tree Curation for Carnitine Pathway Genes

To perform accurate selection tests for 394 genes of interest, sequences were filtered and checked for naming convention discrepancies via a set of custom scripts, and if they did not have matching protein sequences, they were removed from the dataset. Protein alignments and Refseq transcript files were then run in the PERL script PAL2NALto generate protein-informed codon alignments ^31^. Codon alignments were subject to further scrutiny and edited with a custom cleanup pipeline which: 1) Removes stop codons and all sequence downstream of stop codons, 2) Begins the alignment with a codon that is present for at least 70% of species, and 3) Removes sequences (species) that are >50% gaps. After filtering steps 378 gene alignments remained, and a phylogenetic tree was generated for all species included in those alignments (206 species) using TimeTree.org ^32^. Species missing from TimeTree, but included in our sequence data, were added manually, resulting in a 206-species master topology (Supplementary File 1). As each codon alignment requires a corresponding species tree to run selection analyses, a custom python script was used to generate a tree file for each codon alignment, pruned from the master topology using only the species in each alignment file. A full pipeline with all associated scripts is available on Github (https://github.com/drabe004/BearProject_Scripts).

### Tests of Selection

To test all genes for convergent positive selection in hibernating species, sequence alignments and trees were used as input for the HyPhy suite test BUSTED (Branch-Site Unrestricted Statistical Test for Episodic Diversification), which identifies branches in a phylogenetic tree that have experienced episodic positive selection at specific sites across a gene. Briefly, BUSTED uses a codon-based model to compare the fit of a null model, where no sites are under positive selection on any branches, to an alternative model, where some sites may experience episodic positive selection (dN/dS > 1). First, to check for signal of positive selection in hibernating species only, a list of hibernating species (Supplementary File 2) was used to assign tree tips to a “foreground” with remaining tips assigned to “background.” A custom R script assigned internal branches to the Foreground, using simple maximum parsimony in the Castor package ^33,34^. Second, to check for signal of pervasive selection across the tree, a second round of tests was run designating no branches to the foreground (hereafter referred to as “hypothesis free” test). Lastly, to verify that the main source of signal for selection was attributable to hibernating species, a drop test was performed, in which we removed all foreground species from the tree/alignments, and subsequently ran a hypothesis-free BUSTED test ^35,36^. To test for positive selection in hibernating bears only (from which our physiological signal was derived), we also ran tests of selection designating bears (genus *Ursus*) only as foreground taxa. All BUSTED tests were run using the “-srv Yes” flag to account for synonymous rate variation. Results of tests were assessed using a likelihood ratio test, and tests with bears and hibernators in the foreground were compared to hypothesis-free tests and drop tests to compile a list of genes under convergent positive selection which are robust to type 1 error caused by selection in hibernators and non-hibernators.

To test for signals of relaxed selection, we used the RELAX test implemented in the HyPhy suite^37^. Briefly, this method compares the fit of models in which both negatively selected sites and positively selected sites are trending towards an omega value of 1 (neutrality), to a more standard model of positive selection where omega is increasing across all sites. We use the same foreground and background tests as in BUSTED tests, with the exception that RELAX does not allow for a “hypothesis free” option.

Because bats made up a large portion of the hibernating taxa, there was a risk that genes associated with other bat physiologies unrelated to hibernation would bias the dataset by driving signal for selection disproportionately. To account for this, we also reran BUSTED and RELAX (including drop tests) with only a single (hibernating) bat representative (*Myotis myotis*). A table outlining each nested test, including the dataset, foreground species, and interpretation is available in Supplementary Table 2.

### Alignment and Gene Tree Curation for RER Converge

Trees for 120 mammal species were generated from protein sequences downloaded from ^38^. When multiple versions of a protein were included, only the version with the greatest number of species was retained. All “*”, “?”, and “X” symbols were converted to gaps (“-”), species in which the gene was missing (i.e. all “-”) were removed, proteins with fewer than three species were removed, and protein sequences were aligned using the msa package in R ^39^. After aligning sequences, the RERconverge “estimatePhangornTreeAll” function ^40^ with default parameters was used to calculate branch lengths. Briefly, the function uses phangorn ^41^ functions to generate maximum likelihood branch lengths assuming a supplied topology (Supplementary File 3) using the LG model with k=4. The supplied topology was based on one provided with the alignments and adjusted to match the high-confidence tree originally reported in ^42^ based on ^43^ and ^44^. After generating trees, the RERconverge readTrees function was used with default parameters to convert the trees to the proper format for further analyses and generate a master tree with branch lengths representing average evolutionary rate across many regions. In total, we produced 19610 gene trees across 120 mammalian and used these as input for convergent rate analysis.

### RERconverge

To identify genes with convergent evolutionary rates in hibernating mammals we ran RERconverge with 19610 gene trees derived from 120 mammalian genomes that included 14 hibernating species (Supplementary Table 3). Convergent rate analysis is a method of examining whole-genome data by transforming single-copy gene trees to trees where branches represent relative evolutionary rates of protein evolution, constrained by a known species topology. Foreground species which have a common, convergently evolved phenotype (e.g. hibernation) are noted and RER converge identifies genes which are evolving with slower or faster relative evolutionary rates when compared to background species (e.g. non-hibernators). For these data, we used the 14 hibernating species as foreground taxa. All analyses were run using the “permulations” (permutated simulations) option, which evaluates the significance of evolutionary rate shifts by permuting trait values across the phylogenetic tree and recalculating relative evolutionary rates to create a null distribution for comparison. We used Benjamin-Hochberg method to correct for multiple test correction in R.

Similar to selection analyses, the over-abundance of bats representing hibernators was considered as a potential source of bias. To explore the influence of this bias on the dataset, the following nested RERconverge analyses were run: 1) Removing bats from the dataset entirely and 2) Setting bats as the only foreground. We report the RERConverge results for all hibernators alongside the associated significance data for the same dataset using only bats as foreground species and report the rank of each gene by adjusted p-value. This allows the interpretation of this data to be viewed unaltered but within a context of how strongly each gene might be influenced by signal from the bat lineages. This methodology improves on past work which has used Bayes factor analysis to account for overabundance of bat signal, as it allows us to see convergent rates of hibernating genes with a continuous weighted measure of the influence of one lineage without having to assign cut-off values ^45^.

Genes significant in RERconverge analysis were further tested for positive and relaxed selection. Because RERconverge data is derived from protein alignments, not DNA, each significant RER gene (30) was checked for availability on NCBI ortholog database and, if available, were downloaded and filtered as detailed above (the same as KEGG pathway genes). These genes (27 total available) were also tested for positive and relaxed selection using the same methods detailed above.

### Enrichment Analysis

To assess whether genes identified as significant (either by RERconverge or HyPhy tests of selection) were significantly enriched for specific function, enrichment analyses were performed on all datasets using the DAVID bioinformatics resource (https://david.ncifcrf.gov/). Because genes under selection would already be enriched for certain pathways by virtue of being chosen for their association with carnitine metabolism, and only a small number of genes were recovered both under selection and as evolving under convergent rates (RERconverge), we used the following approach to reduce statistical noise and dataset bias. Each dataset (19610 genes for RERconverge genes and 378 genes associated with carnitine metabolism) was tested for enrichment by comparing it to all annotated human genes in DAVID. Subsequently, a subset of genes identified as significant in RERconverge analyses, positive selection analyses, and relaxed selection analyses (separately), were similarly tested for enrichment by comparing to all human genes in DAVID. Enrichment terms reported as significant in both the total set and the subset were removed, as enrichment of these terms would represent dataset-specific bias. Terms that were significantly enriched in the subsets only were kept and used to generate figures and final enrichment tables (Supplementary Figure 2).

## Results

### Bear Blood Metabolic Data

To identify clear contrasts in 159 metabolic indicators, we focused on comparisons of early and late denning. Classification performance of the sPLS-DA model between early and late denning showed that these two components reflect 23% of the original data’s variation (component 1: 14% of the variance, component 2: 9%) (Figure 2A, Supplementary Data 1).

The sPLS-DA model identified three metabolic indicators which clearly defined the early denning cluster: carnitine esterification, beta oxidation, and ratio of acetylcarnitine to carnitine (Figure 2B, Supplementary Data 2). Of the raw metabolites that are included in metabolic indicator calculation, carnitine and acetylcarnitine, both are significantly decreased during early denning (Figure 2D, Supplementary Figure 1). All three indicators showed an inverse relationship with raw carnitine abundance (and to a lesser degree acetylcarnitine), as the metabolic indicators were all significantly higher in early denning, while raw carnitine was significantly lower in early denning compared to all other time points (Figure 2C&D, Supplementary Figure 1). These findings suggest that metabolic pathways related to carnitine (including fatty acid metabolism and beta-oxidation) are critical to metabolic shifts from the active period to hibernation. Because pathways altering carnitine levels may be critical to hibernation as well as important potential therapy targets, our subsequent bioinformatic analysis focused on pathways which were related to carnitine in KEGG.

### C*onvergent Rate Analysis*

Phylogenetic analysis of 120 mammalian genomes resulted in a total of 19,610 gene trees (Supplementary File 4). Convergent rate analysis selecting all hibernators as the foreground identified several genes evolving convergently at significantly slower rates (negative Rho) in all hibernators when compared to non-hibernating mammals (Table 1). Several genes were also identified as evolving at significantly faster rates convergently when compared to non-hibernating species (Table 1). No genes were identified as significant when bats were removed from the foreground, suggesting that convergence with bats genes influence these signals. When bats were placed as the only foreground species, 573 genes were identified as evolving at significantly convergent rates. However, genes identified under convergent rate evolution for all hibernators were not a simple subset of the most significant bat-convergent genes, but rather were distributed across the 573 genes ranked by their p value (Table 1). This ranking system reveals that while the significance of RER analysis for some genes is clearly mainly driven by bat lineages, (e.g. GJA8, NDST3, CNGB3), others are more so driven by the convergence of bats and other hibernators (Table 1). Using an extensive literature review, we categorized each gene by the hibernation-related phenotypes they are most likely to contribute to. We found genes evolving at convergent rate in hibernators related to tissue regeneration and repair (5 faster, 1 slower), gene regulation (2 faster, 3 slower), metabolic shifts (4 faster, 1 slower), oxidative stress and hypoxia (3 faster, 1 slower), coagulation and dehydration (2 slower), skeletal muscle preservation (2 faster, 1 slower), thermogenesis and cold tolerance (1 faster), and innate immunity (1 slower) (Figure 3, Supplementary File 5).

**Figure 3.**
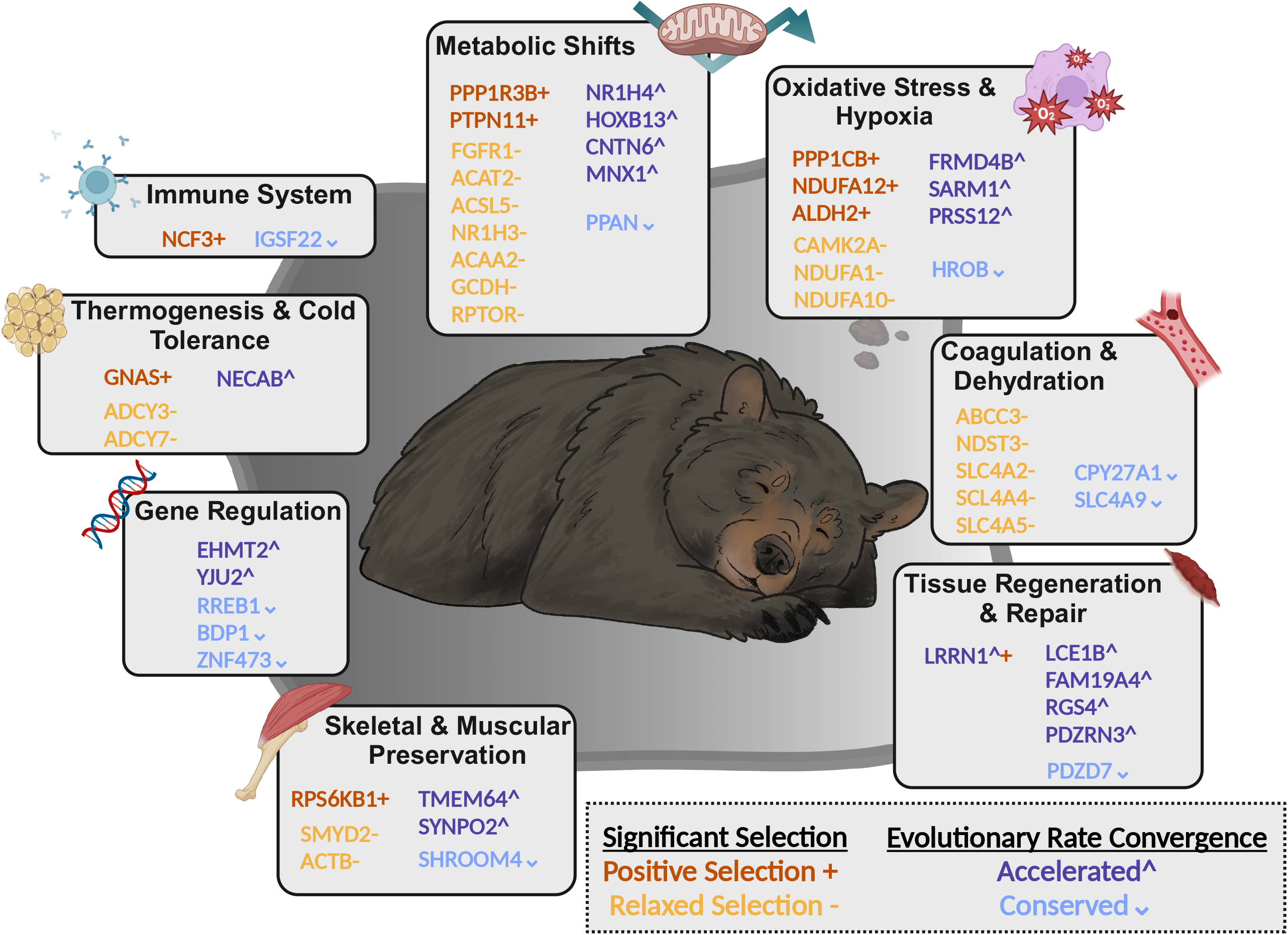
Summary of genes found to be significant for tests of positive selection (+), tests of relaxed selection (-), convergently faster rates of evolution (^), and convergently slower rates of evolution (v) among hibernators. Details of each result can be found in supplementary table 5, and an expanded discussion of each can be found in supplementary file 5.

**Table 1.** Results of convergent rate analysis performed with RERconverge binary trait analysis placing all hibernating mammals in the foreground, using 19610 gene trees derived from a 120-mammalian genome (protein) alignment. Genes evolving under significantly convergent evolutionary rates using ‘permulations’ (phylogenetically informed simulations) and adjusted for multiple test correction are reported below. Significance for the same gene when run using only bats as foreground taxa is also reported, as well as a rank number which represents where the gene (significant for all hibernators) fell in a distribution of bat-only significant genes ordered by p-value. For example, when bats only are in the foreground CNGB3 was the number 1 most significant gene, whereas ZNF473 was the 303rd. Asterisks denote genes which were also significant in tests of selection (See Supplementary Tables 5 and 6). Rho is a measure of correlation between the relative evolutionary rate and the trait over all branches (negative numbers are slower than average and positive numbers are faster than average). N is the total number of branches in the gene tree, and p.adj values are p values after correction for multiple tests using the Benjamini-Hochberg method.

| Genes Evolving at Convergent rates across all Hibernators (RER Converge) |  |  |  |  |  |
| --- | --- | --- | --- | --- | --- |
| Genes Evolving Slower than Average |  |  |  |  |  |
| Gene Symbol | Rho | N | p.adj | p. adj Bats only | Rank (1-573) |
| IGSF22 | -0.230058392 | 201 | 0.053724143 | NS | NS |
| ZNF473 | -0.218498392 | 221 | 0.054110144 | 0.031039 | 303 |
| SLC4A9 | -0.225346488 | 210 | 0.053724143 | 0.026846 | 275 |
| CYP27A1 | -0.262287744 | 195 | 0.03084742 | 0.01288 | 107 |
| PPAN | -0.233693402 | 192 | 0.054110144 | 0.01288 | 111 |
| HROB | -0.225983985 | 221 | 0.050117271 | 0.011678 | 92 |
| PDZD7 | -0.237011544 | 223 | 0.038115889 | 0.010806 | 84 |
| RREB1 | -0.226448056 | 213 | 0.051857967 | 0.00923 | 68 |
| BDP1 | -0.223987666 | 228 | 0.050117271 | 0.006494 | 45 |
| SHROOM4 | -0.228902052 | 212 | 0.051397465 | 0.004722 | 34 |
| Genes Evolving Faster than Average |  |  |  |  |  |
| Gene Symbol | Rho | N | p.adj | P. adj Bats only | Rank (1-573) |
| FAM19A4 | 0.365000536 | 95 | 0.038115889 | NS | NS |
| PDZRN3 | 0.252750446 | 169 | 0.053724143 | 0.045897 | 452 |
| MNX1 | 0.293421504 | 167 | 0.018624689 | 0.024597 | 238 |
| FRMD4B | 0.25743257 | 187 | 0.038115889 | 0.021874 | 191 |
| LCE1B | 0.294470449 | 127 | 0.051857967 | 0.020725 | 184 |
| PRSS12 | 0.243972247 | 201 | 0.045888219 | 0.017695 | 163 |
| LRRN1 | 0.273312637 | 158 | 0.046117471 | 0.013337 | 117 |
| SARM1 | 0.260834114 | 175 | 0.045888219 | 0.013054 | 113 |
| RGS4 | 0.245855993 | 173 | 0.054110144 | 0.010488 | 80 |
| NECAB1 | 0.380097581 | 75 | 0.053724143 | 0.00923 | 65 |
| TMEM64 | 0.258961508 | 163 | 0.051857967 | 0.008652 | 59 |
| YJU2 | 0.242320316 | 179 | 0.054110144 | 0.008129 | 54 |
| HOXB13 | 0.272711205 | 153 | 0.050117271 | 0.004722 | 28 |
| CNTN6 | 0.398099728 | 120 | 0.002248326 | 0.004722 | 24 |
| SYNPO2 | 0.273467722 | 207 | 0.01031322 | 0.004722 | 17 |
| CX3CL1 | 0.231588736 | 211 | 0.050117271 | 0.003785 | 15 |
| GJA8 | 0.284920435 | 195 | 0.01031322 | 0.002137 | 6 |
| NDST3* | 0.262815157 | 162 | 0.051397465 | 0.002026 | 3 |
| CNGB3 | 0.243104716 | 210 | 0.038115889 | 0.002026 | 1 |

### Positive Selection in Genes Associated with Carnitine Metabolism

To assess signal of positive selection across 378 genes identified as potential contributors to carnitine metabolism (plus 27 genes identified as significant in RER analysis for a total of 405), we performed several nested statistical tests of selection using BUSTED in the HyPhy software suite. Grouping species into a single foreground can be a useful way to identify convergent selection, however, it also runs the risk of elevated type three error, or misattribution of the origin of signal for selection ^35,36^. We, therefore, report results for all selection tests in the context of a set of nested tests designed to narrow down convergent signatures of positive selection. A total of eighteen genes were found to be evolving under positive selection (Figure 3, Supplementary Table 4). When all but one hibernating bat is removed from the dataset, half of these genes remain significant (Supplementary Table 5). Though six of these remaining nine genes are also significant in a hypothesis free test (with no foreground specified), all of them lose significance when performing a drop test (removing all hibernators from the dataset, and performing a hypothesis free test), indicating signal of positive selection originates primarily from the lineages of hibernators, and not from bats alone (Supplementary Table 4). We did a similarly comprehensive literature review of genes under positive selection (which retained significance when the majority of bats were removed Supplementary Table 4), and found genes under positive selection in hibernators were related to oxidative stress and hypoxia (3), metabolic shifts (2), thermogenesis and cold tolerance (1), and innate immunity (1) (Figure 3, Supplementary File 4).

### Relaxed Selection in Genes Associated with Carnitine Metabolism

Statistical signal of increased diversifying mutations on a gene may be indicative of positive selection or relaxation of selection, and telling apart these two processes is notoriously difficult ^37^. To distinguish these two scenarios, we use the test RELAX in the HyPhy suite which tests the fit of a model in which all sites’ omega values are either moving towards 1 (neutrality) or away from 1^37^. Similar to tests of positive selection, signal of relaxed selection may be subject to misattribution (type three error), and so we report the results of nested hypothesis free (BUSTED) tests as well as the same tests run removing bats entirely from the dataset (Figure 3, Supplementary Table 5). Six genes were found to be evolving under relaxed selection in all hibernators and remained significant when removing bats from the data. An additional 14 genes were significant for relaxed selection when only bears (*Ursus spp*) were placed in the foreground (Figure 3, Supplementary Table 5). All but two genes (of 20 total) were not significant in a hypothesis free BUSTED analysis, indicating a potential for high rates of diversification across the tree in these 18 genes (Figure 3, Supplementary Table 5). We also placed these genes under hibernation-related phenotypic categories based on an extensive literature review, and found genes related to metabolic shifts (7), coagulation and dehydration (5), oxidative stress and hypoxia (3), skeletal and muscle preservation (2), and thermogenesis and cold tolerance (2) (Figure 3, Supplementary File 5).

### Enrichment Analysis

To identify annotated gene function which may be overly represented in genes identified by RER converge, or HyPhy selection tests, we used the DAVID bioinformatics resource to group and test functional enrichment ^46^. No genes identified in the convergent rate analysis (30) were over-represented for any functional annotation term when using either the full 19610 genes derived from 120 mammalian genomes as background or the default DAVID human genome database (all human annotations). Genes experiencing both positive and relaxed selection (38) were significantly enriched for several terms when using all human annotations as a background. Terms that were enriched uniquely in the 38 genes experiencing positive or relaxed selection and not when all (378) carnitine-associated genes where used in the same analysis are reported in Supplementary Figure 2 (full data in Supplementary File 6). Terms that were strongly enriched in genes under positive selection are strongly linked to hibernation-related phenotypes including: Thermogenesis, Respiratory Chain, NADH Dehydrogenation, and Mitochondrion. Genes under relaxed selection were enriched for a larger number of terms with less clear links to hibernation, with a notable exception for terms related to insulin and glucose metabolism (Supplementary Figure 2).

## Discussion

Though carnitine has been identified as a key metabolite in hibernation previously in squirrels, it has not been specifically indicated in bear hibernation ^47–53^. However, recent work on brown bears (*Ursus arctos*) has shown that during hibernation, there is a metabolic shift that reduces the production of Trimethylamine (TMA, a byproduct of carnitine metabolism by gut bacteria) and trimethylamine N-oxide (TMAO), whereby bears’ metabolisms favor the production of betaine over TMAO ^20^. Our data show that carnitine levels are reduced during black bear (*Ursus americanus*) hibernation, which may facilitate the metabolic shift seen in recent work and may be the ultimate driver of shifts to betaine production. Understanding the pathways and genes that contribute to these shifts help to reveal the genetic basis of this vital adaptation as well as shed light into avenue of carnitine regulation with respect to its role as a widely used biomarker for a diverse range of conditions ^54^.

By integrating metabolomics and comparative genomics, we identified genes evolving convergently across diverse lineages of hibernating mammals that may underlie critical phenotypic changes unique to hibernation, particularly with respect to metabolic shifts and carnitine metabolism (Figure 4). Though many genes have previously been identified as having altered regulation during hibernation, presence of signal from coding sequences suggests that these genes are also under diverse selection pressures (positive, relaxed, or negative) to alter, lose, or maintain their function at the sequence level. We also identify several novel genes, previously unknown to contribute to hibernation but related to several key hibernation-related physiologies.

Below we highlight significant genes in this study related to metabolic shifts and cover a more extensive review of significant genes related to other aspects of hibernation physiology in the Supplementary Materials (Figure 3, Supplementary File 5). Enrichment analysis showed that genes under positive selection are significantly enriched for fewer terms than genes under relaxed selection, which is unsurprising given that relaxation of selection is expected to be a more neutral process (Supplementary Figure 3). However, in our own literature review we find genes under relaxed selection to be directly related to metabolic shifts and carnitine, with several known to cause beneficial effects when inhibited or knocked out in mice. While carnitine-related genes under selection are more heavily focused on metabolism and oxidative stress related functions, convergent rate analysis reveal genes that are more diverse in function (such as regulatory elements and immune system genes) and may contribute to more diverse aspects of the hibernation physiology (Figure 3, Supplementary File 5). Both approaches reveal important insights about the mechanisms that underlie shifts in metabolic indicators found in this study, as well as other aspects of hibernation physiologies found in previous studies in a wide range of hibernating species. Below, we use the following shorthand symbols to discuss genes under positive selection: ***GENE+***, relaxed selection: ***GENE-***, faster convergent evolutionary rates: ***GENE^***, and slower convergent evolutionary rates: ***GENE***_***V***_ in hibernating mammals (Figure 3).

With carnitine closely linked to fat metabolism, it is unsurprising that the greatest number of genes (14) recovered in our dataset are related to metabolic shifts (Figure 3). ***PPP1R3B+*** encodes a glycogen-targeting subunit of protein phosphatase 1 (PP1) that plays a crucial role in regulating hepatic glycogen synthesis, storage, and fasting glucose metabolism. PP1 has been shown to be downregulated in the liver and brain of torpid squirrels and hibernating Syrian hamsters and has been uniquely shown to be unaffected by temperature, though mechanisms for these changes are unknown ^55,56^. While previous work showed that profound inhibition/downregulation and temperature stability of this pathway is vital for processes involved in hibernation, our work identifies *PPP1R3B* as a key target gene under positive selection which may be contributing to these functional shifts ^56^.

With respect to hibernation, much interest has been paid to the function of ***FGFR1-***as a receptor that mediates the effects of *FGF21* in coordination with the co-receptor β-Klotho, which can induce a torpor like state in mice, is upregulated in hibernating mammals, and is linked to the regulation of food intake, body weight, and DIO2 (a thyroid hormone enzyme) expression ^57–59^. The *FGFR1* signaling pathways regulates glucose homeostasis, lipid metabolism, and thermogenesis, and inhibition of the *FGFR1* signaling pathways has been shown to reduce adiposity, body weight, and increase insulin sensitivity ^22,59,60^. Recovery of signal of relaxed selection in this gene may indicate a beneficial effect of a partially functional *FGFR1*, like what is seen when this protein is inhibited in mice.

Two genes identified as being under relaxed selection (***ACAT2-*** and ***ACSL5-***) are involved in lipid metabolism and have previously been identified as being differentially expressed in several studies of hibernating mammals and overwintering frogs ^61–64^. *ACSL5* activates long-chain fatty acids for biosynthesis, while *ACAT2* is primarily involved in cholesterol metabolism and transport, its overexpression in mice may be protective of high-fat-diet-induced weight gain ^65–68^. Both *ACAT2* and *ACSL5* are directly linked to carnitine as they play key roles in lipid metabolism, particularly in the activation and transport of fatty acids for β-oxidation. ***NR1H3-***has not been previously linked to hibernation but is a nuclear receptor that regulates the expression of genes involved in cholesterol, lipid, and glucose metabolism by promoting the efflux of cholesterol from cells to prevent accumulation as well as influencing inflammatory immune response ^69^. ***ACAA2-*** and ***GCDH-***are both involved in fatty acid β-oxidation, and similarly have been previously identified as being differentially expressed across several diverse mammals as well as in hibernating frogs ^63,70–75^.

Other notable genes involved in lipid metabolism are ***NR1H4^, HOXB13^***, and ***CNTN6^***, all of which we find to be evolving at convergently faster rates among hibernators. *NR1H4*, differentially expressed in hibernating black bears, was recently identified as an upstream regulator of genes that showed altered expression after a mid-hibernation feeding of captive brown bears and was predicted to facilitate metabolic shifts to glucose metabolism ^63,76^. *HOXB13* and *CNTN6* have both been shown experimentally to be strongly associated with fat deposition, fat percentage, and metabolism ^77,78^.

Evolving slower than average in hibernators, ***PPAN***_***V***_ (Peter Pan homolog) is a gene involved in cell growth and proliferation, playing a key role in ribosome biogenesis and protein synthesis, and has been shown to be downregulated during diapause in fruit flies (*Drosophila melanogaster)* ^79^. Ribosome hibernation factors (e.g., RSFS) are known to be employed to reduce energy consumption by silencing ribosomes and keeping them inactive while preserving their readiness for translation ^80^. Slowed evolutionary rates in PPAN may preserve its compatibility and function as a ligand for silencing factors.

In addition to shifting to fat metabolism, hibernating species also cope with prolonged starvation by inducing a state of decreased insulin sensitivity (mimicking type-2 diabetes) and reducing insulin production. Three genes identified in this study, ***PTPN11+, RPTOR-***, and ***MNX1^*** are all critical to these processes. *RPTOR* has been shown to be upregulated and hyperphosphorylated in hibernating mammals and in leeches (respectively), and RPTOR knockout mice were protected against high-fat-diet-induced obesity, insulin resistance, inflammation, and expansion of adiposity in bone marrow ^81–85^. MNX1, not previously identified to be involved in hibernation, is a homeobox gene that encodes a transcription factor essential for the differentiation and proper functioning of insulin-producing beta cells in the pancreas ^86^. *PTPN11* has been linked to glycogenesis and obesity-induced infertility and may facilitate maintenance of fertility despite the massive metabolic changes (weight loss and gain) required to weather a hibernation season ^87–90^.

Genes contributing to oxidative stress and hypoxia also make up a large proportion of genes recovered in this work and emphasize the inter-relatedness of metabolic shifts and hypoxic stress (Figure 3, Supplementary File 5). Specifically, genes encoding subunits of Mitochondrial Complex 1 (NDUFA genes) are novel to hibernation biology and are indicated three separate times in our selection analyses. Genes related to coagulation and dehydration (8), skeletal and muscular preservation (6), tissue representation and repair (5), gene regulation (5), thermogenesis (4), immunity (2), as well as genes that are likely to be important strictly in bat lineages are important findings of this work and are discussed in greater detail elsewhere (Supplementary File 5). Of note are the SCC4 family genes, which regulate pH balance and ion homeostasis, were recovered four separate times in our analyses, and are novel to hibernation biology.

Metabolomic data in this work indicate carnitine metabolism as a significant physiological component of hibernation. This is particularly relevant to critically ill sepsis patients as degree of mitochondrial dysfunction and increased mortality have been linked to early elevated levels of acylcarnitine^91–93^. Measurements of the carnitine pool could lead to important advances in the diagnosis, prognosis, and treatment of these diseases ^54^.

While it is unclear whether lower carnitine levels are a consequence or a cause of metabolic shifts in hibernation, manipulation of carnitine levels and/or pathways genes which regulate carnitine levels discusses here may be a productive avenue of hibernation-based therapy design.

Selection and convergent rate analysis has revealed both known and novel genes that may underlie the diverse physiological adaptations responsible for the hibernation phenotype, and serves to guide research aimed at finding clinically relevant therapy targets. While we aimed to outline how genes may contribute to each physiological component above, it is important to consider that many of these genes are involved in highly pleiotropic pathways. Further work is needed to elucidate the phenotypic consequences of these genes, how they might be linked to hibernation, and how they can be utilized for hibernation-based therapies.

## Acknowledgements

Dr. Iles was issued a permit #35824 by the State of Minnesota Department of Natural Resources and Division of Fish and Wildlife to possess and transport samples of blood and bile and portions or derivatives thereof, including but not limited to enzymes, other bioactive molecules, and genetic materials (hereinafter, “samples”), collected from black bears (*Ursus americanus*) in Minnesota. An Institutional Care and Use Committee Wildlife protocol was approved during the time of sample collection from the University of Minnesota,#1411-32045A. We would like to acknowledge the Minnesota Department of Natural Resources for their permitting and efforts in collecting bear samples in Minnesota for the past decade. We would specifically like to recognize Dr. David Garshelis and Spencer Rettler for their wildlife expertise and continued collaboration for obtaining samples and fieldwork. We would also like to thank Dr. Timothy Laske and Dr. Paul Iaizzo for their field work. Thank you to the Center for Mass Spectrum Analysis and Proteomics for running the Biocrates plasma samples and their support in the analysis. We would like to thank the Chair of the Department of Surgery, Dr. Sayeed Ikramuddin for his departmental and research support.

