## Supplementary for "Genes Underlying Adaptive Physiological Shifts Among Hibernating Mammals": Supplementary Figures 1-2.docx

**Supplementary Figure 1. Bar plots of metabolic indicators and raw metabolites for summer, early denning, and late denning periods. Asterisks indicate significance at alpha = 0.05.**

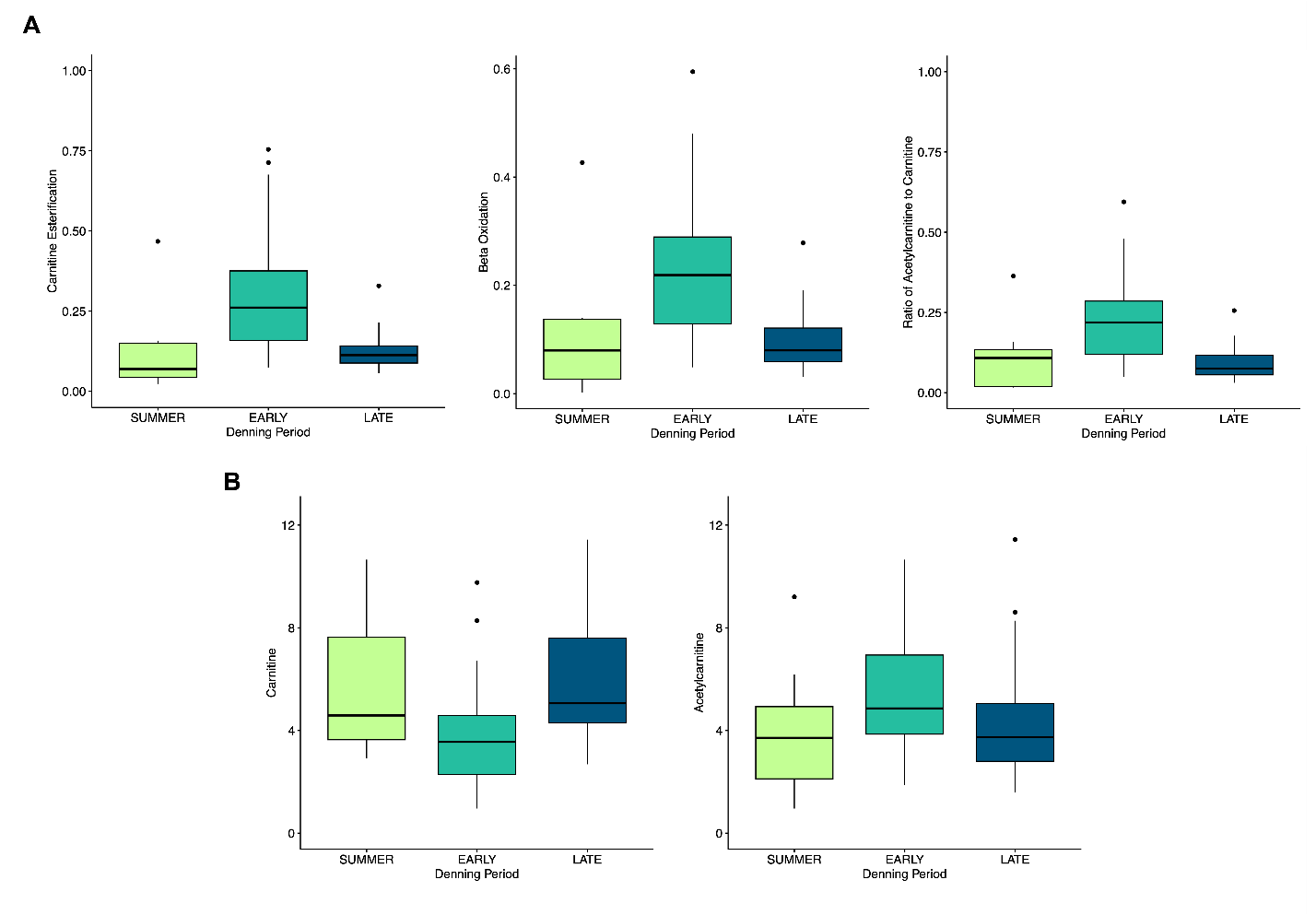

**Supplementary Figure 2.** Enriched annotation terms for genes under selection. The size of each chunk in the stacked chart corresponds to the number of genes contributing to the enrichment of a given annotation. The size is proportional to the number of contributing genes relative to the total number of genes contributing to all annotations (n = 92 genes). Thus, the size of each chunk represents the count for the annotation term divided by the total, and the chunks cumulatively sum to 1. Annotation sources are listed in parentheses following the annotation name.

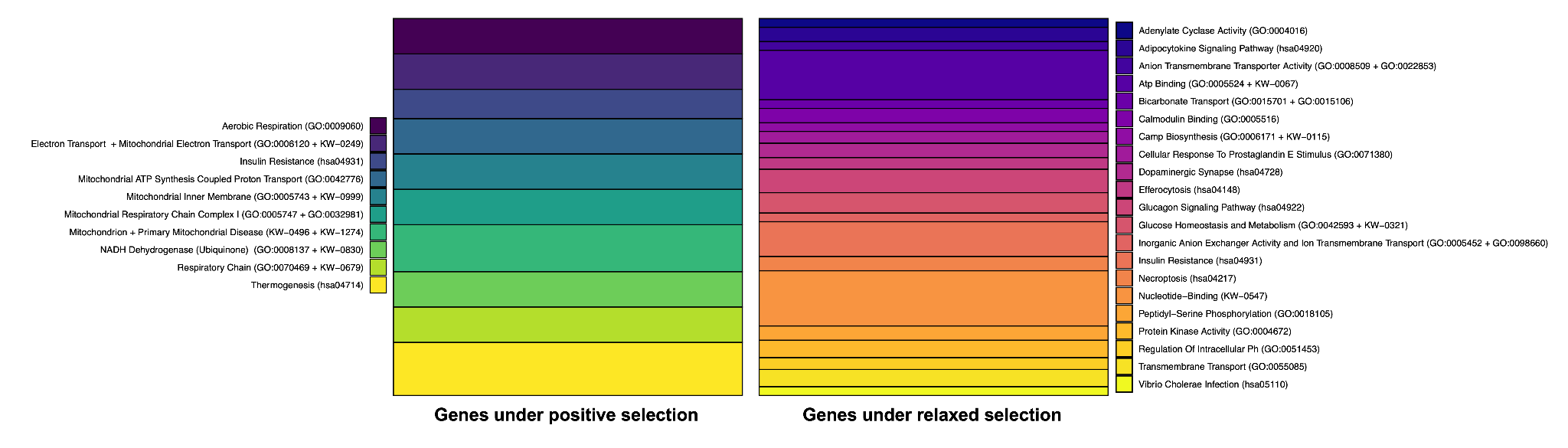
