## Supplementary for "Genes Underlying Adaptive Physiological Shifts Among Hibernating Mammals": Supplementary File 5.docx

**Supplementary Discussion**

***Genes Underlying Oxidative Stress and Hypoxia (9)***

Several genes revealed in this study appear to be related to oxidative stress and hypoxia, which are hallmarks of the extreme metabolic shifts during hibernation. ***PPP1CB+*** is a protein phosphatase involved in cell division and metabolism, is upregulated under hypoxia in Tibetan chickens ^1^. Though this gene has not previously been identified as a candidate for hypoxia tolerance in mammals, positive selection on *PPP1CB* across hibernators indicates a role in managing metabolic suppression during hypoxia.

***CAMK2A-*** encodes a subunit of the calcium/calmodulin-dependent protein kinase II (CAMKII) that plays a crucial role in regulating calcium signaling. However, reactive oxygen species (ROS) can activate the kinase independently of calcium leading to dysregulation, and ROS-insensitive CAMKII has been shown to be protective of host of diseases related to oxidative stress ^2^. This cost of increased oxidative stress has been cited as an example of age-related antagonistic pleiotropy, and suggests that relaxation of this gene in hibernating species may facilitate enhanced oxidative stress tolerance. Inhibition of CAMK2 has also been shown to improve liver insulin signaling, insulin resistance, skeletal muscle INSR signaling, and treatment of obesity related glucose intolerance ^3,4^.

***NDUFA12+***, ***NDUFA1-*,** and ***NDUFA10-*** all encode subunits of mitochondrial complex I, which transfers electrons from NADH to ubiquinone and supports ATP production in cells ^5^. Complex I is a major source of reactive oxygen species (ROS) under hypoxia, and reduced glucose oxidation via hypoxia-inducible factor pathways (HIFs) decreases electron flow through Complex I, conserving oxygen, minimizing ROS production, and reducing reliance on Complex I for ATP generation, thereby promoting hypoxia tolerance ^6^. Previous work has identified Complex 1 genes as differentially expressed during hibernation in both ground squirrels and black bears (*NDUFA12* and *NDUFA 10*, respectively)^7,8^. Signatures of positive (*NDUFA12*) and relaxed (*NDUFA10*, *NDUFA1*) selection in these genes suggests that hibernators may be evolving variants which are either more efficient or more easily inhibited by HIFs under hypoxic conditions.

Another gene involved in mitochondrial function that we find to be under positive selection is ***ALDH2+***, a mitochondrial enzyme that detoxifies aldehydes, which is linked to increased risk of oxidative stress-related diseases. *ALDH2* expression has been shown to be lower in ischemia-reperfusion injury models but is increased during the torpor phase in hibernating thirteen-lined ground squirrels ^9–11^. In the northern red-backed vole (*Myodes rutilus*), a species that remains active during extreme winter conditions, activity of *ALDH2* (along with other aldehyde dehydrogenase enzymes) in the liver significantly decreases and is associated with a winter increase in blood acetaldehyde concentration ^12^. This suggests *ALDH2* may play a protective role in hibernation by supporting enhanced fatty acid metabolism and guarding against ischemia-reperfusion injury.

***HROB⌄*** promotes DNA repair through homologous recombination, likely contributing to enhanced DNA repair mechanisms in response to oxidative stress is (a well-established aspect of the hibernating phenotype) ^13–15^.

Several genes identified as evolving at a convergent accelerated rate among hibernators have functions related to neuroprotective mechanisms. Previous work has shown hyperphosphorylation of tau proteins during hibernation which resembles the pathological changes seen in neurodegenerative diseases but also stabilizes the cytoskeleton and protects neurons during the low-energy state of hibernation. **FRMD4B^** may be important for hibernation by modulating tau secretion through the FRMD4A-cytohesin-Arf6 signaling pathway, helping to prevent harmful tau accumulation during metabolic suppression, thereby preserving neuronal integrity and preventing neurodegenerative-like damage ^16–18^. ***SARM1^***, responsible for axonal degeneration via NAD+ breakdown, is an established pharmaceutical target for neuroprotection, and may help regulate neuronal survival during hibernation. The connection between tau hyperphosphorylation and *SARM1* may lie in their complementary roles in managing neuronal stress: tau phosphorylation helps to stabilize neurons during hibernation, while *SARM1* could be fine-tuned to regulate axonal survival and degeneration, ensuring that neurons are protected but also ready to recover normal function when hibernation ends ^19–21^. Similarly, ***PRSS12^*** (neurotrypsin) encodes a serine protease which supports nerve repair under metabolic stress ^22^.

***Coagulation and Dehydration (7)***

Prolonged dehydration causes electrolyte imbalances and reduced plasma volume, leading to coagulopathy by disrupting clotting pathways and increasing blood viscosity, which together promote both hypercoagulability and impaired clotting. Several genes revealed in this study are implicated in functions that may contribute to these interlinked physiologies.

Two genes, ***ABCC3-*** and ***CYP27A1⌄***, are involved in mitigating bile acid toxicity and likely mediate mechanisms that prevent the harmful effects of toxin build up during hypometabolism and anuria. *ABCC3* is involved in the efflux of organic anions and drugs from cells which is critical for detoxification and efflux of metabolic waste products, and is differentially expressed/hyperphosphorylated in several hibernating animals^8,23–25^. Suppression, or reduced function of *ABCC3* in hibernators could be beneficial as it may help retain bile acids, such as ursodeoxycholate (UDC), within the liver and biliary tree rather than transporting them out of the liver and into circulation. *CYP27A1* is primarily involved in the conversion of cholesterol into bile acids, which is crucial for the elimination of cholesterol from the body. *CYP27A1* has been shown to be downregulated during hibernation in black bears, likely contributing to reduced cholesterol catabolism and elevated serum cholesterol levels (bears)^26–28^.

***NDST3-*,** a type of glycosaminoglycan, has been shown to be one of the three most significantly differentially expressed genes during hibernation in black bears(), and its expression is upregulated specifically in the kidney^8^. Glycosaminoglycan are critical components of selective vascular filtration and are known to be involved in water conservation during torpor in turtles^29^. It is also involved in modifying clotting factor inhibitors and may more generally be involved in the protection against coagulopathy ^30^.

The *SLC4A* genes belong to the SLC4 family of solute carriers, which encode a group of bicarbonate transporters involved in maintaining pH balance and ion homeostasis in cells ^31^. ***SLC4A2-*** (anion exchanger 2), ***SLC4A4-*** (sodium bicarbonate cotransporter NBCE1), and ***SLC4A5-*** (sodium bicarbonate cotransporter NBCE2), were all recovered as genes showing signatures of relaxed selection in hibernating mammals. Signal for relaxed selection among these genes may indicate that hibernators are shifting regulation of electrolyte balance to other pathways to cope with extreme dehydration and anuria. In addition to finding several SLC4A proteins under relaxed selection, we also uncover ***SLC4A9⌄*** as evolving relatively slower in hibernating mammals. This may indicate a shift in genes being used primarily to regulate pH balance in hibernators (from *SLC4A2,4*, and *5* to *SLCA9*). It might also indicate physiological constraints where changes in pH regulation and bicarbonate reabsorption are not tolerated, as *SLC4A9* is expressed primarily in the saliva, pancreas, and airways ^31,32^. Given how many times this gene family arises in this dataset and its lack of prior mention in hibernation studies, future research is merited to decipher its role in the hibernating phenotype.

***Skeletal and Muscular Preservation (6)***

Despite long term disuse, hibernating mammals do not experience the significant bone and muscle degradation seen in during human immobility^27,33–36^. Molecular mechanisms that facilitate resistance to osteoporosis and muscle wasting are of broad interest, yet little is known about the genes that underlie these functions. In this study we find two genes ***RPS6KB1+***, and ***SMYD2-*** under positive and relaxed selection, respectively, that are linked to muscular preservation and resistance to atrophy. *RPS6KB1* encodes a serine/threonine kinase involved in the mTOR signaling pathway. *RPS6KB1* has been shown to be upregulated during hibernation in the muscle tissue of thirteen-lined ground squirrels and black bears, and it is thought to be a key mechanism of avoiding muscle disuse atrophy^11,37,38^.

*SMYD2* plays a key role is in regulating transcription, and has been previously identified as a key protein in muscle atrophy prevention in two species of hibernating squirrels, and is also significantly reduced during diapause in frogs^39–42^. *SMYD2* has also been shown to play a protective role in stabilizing titin (elastic protein found in the sarcomeres of muscle cells) during hibernation, aiding in muscle maintenance and preventing protein degradation in the ground squirrel's skeletal muscles^43^.

***SHROOM4⌄*** and ***ACTB-***, are both involved in the actin cytoskeletal function as SHROOM4 primarily regulates actin organization, whereas *ACTB* forms the structural backbone of actin filaments^44–46^. *ACTB* is generally considered to be a housekeeping gene and has been explicitly used as control genes in several studies examining differential expression in hibernation ^27,47,48^. However, *ACTB* has been found to be significantly under-expressed and inhibited by microRNAs during hibernation in arctic ground squirrels (*Spermophilus parryii*)^49,50^ .

***TMEM64^*** regulates calcium signaling during osteoclast differentiation by modulating *SERCA2* activity, impacting bone resorption and homeostasis, while also playing roles in mitochondrial ROS production and lipid metabolism in adipocytes^51^. Knockouts studies and survey of osteoporotic patients show that there is a strong relationship with reduction of expression of ***TMEM64^*** and osteoporosis^52^. Though this gene has not previously been identified as one that is especially important to hibernators, it may be an important mechanism of bone mass regulation in hibernators.

***SYNOP2^*** is essential for smooth muscle function, regulating actin polymerization and maintaining muscle contraction, with reduced expression impairing differentiation, particularly in vascular injury^53^. Furthermore, a non-coding RNA within the *SYNOP2* intron (*SYISL*), is involved in regulating muscle development and promotes muscle atrophy ()^54^. Accelerated evolution on this gene may indicate an adaptive mechanism to prevent atrophy during hibernation.

***Tissue Regeneration and Repair (5)***

Prolonged inactivity directs stress toward a variety of tissues and is known to cause complications related to tissue damage^55–59^. In this work, we found several tissue-specific genes under convergent accelerated evolution in hibernating mammals that may contribute to tissue resilience during long-term inactivity. ***LCE1B^*** and ***FAM19A4^*** are involved in in skin barrier formation inflammatory response to tissue damage^60,61^. *RGS4* is (along with *FAM194A*) is also involved in brain injury and neuroinflammation. ***RGS4^*** has been shown to be significantly upregulated in the hypothalamus of shrews undergoing Dehnel’s phenomenon, the seasonal shrinking of brain tissue during overwintering , and was upregulated in response to mild traumatic brain injury contributing to a hypometabolic hibernation-like neuroprotective state by reducing glutamate signaling, inhibiting MAP kinase activity, and increasing resistance to calcium-mediated toxicity (^62,63^). ***LRRN1^*** is implicated in neuronal development, axon guidance, synapse formation, and neurite outgrowth^64^. Other tissue specific genes are involved in cardiac structure (***PDZRN3^***) and scaffolding of retinal photoreceptors and inner ear stereocilia **(*PDZD7⌄***) ^65–68^.

***Gene Regulation (5)***

While targeted examination of selection in genes associated with carnitine metabolism yielded several genes specific to known functional physiologies, convergent rate analysis (agnostic of gene function) revealed several regulatory elements which may be critical to hibernation and may also be, more generally, candidates that facilitate physiological plasticity to extreme conditions. ***RREB1⌄***, ***BDP1⌄***, and ***ZNF473⌄*** were all identified as evolving slower in hibernating species and are involved in the regulation of transcription^69,70^. *RREB1* is a zinc finger transcription factor that binds to Ras-responsive elements and has been previously identified as a potential regulatory element responsible for changing expression of PTL, an enzyme which is thought to be responsible for changes in lipid metabolism during hibernation in thirteen-lined ground squirrels^71,72^. ***EHMT2^*** and ***YJU2^*** are both evolving faster in hibernators and have likely roles in epigenetic modification and mRNA splicing (respectively)^73,74^. *EHMT2* is a methyltransferase whose increased expression has been shown in hibernating black bears and plays a crucial role in anoxia tolerance in turtles (with a similar methyltransferase indicated in freeze tolerance in frogs^8,74,75^.

***Thermogenesis and Cold Tolerance (4)***

We recovered several genes which are closely link to and likely candidates for adaptations to cold tolerance. ***GNAS+*** encodes a G-protein subunit that activates adenylate cyclase, increasing cAMP levels and regulating metabolism and energy through hormone production in the endocrine system (thyroid, pituitary, gonads, and adrenal glands)^76^. ***GNAS+*** is highly expressed in the hypothalamus and brown adipose tissue (BAT) of the thirteen-lined ground squirrel throughout hibernation, and is differentially expressed in matrilines, acting as a maternally-expressed promoter and a paternally-expressed inhibitor of non-shivering thermogenesis^15,15,25^. *GNAS* is critical for signaling in preoptic neurons that regulate adaptive thermogenesis, and *GNAS* knockdowns in preoptic neurons in mice significantly reduce nocturnal locomotion, potentially due to metabolic shifts or disruption of circadian rhythms^77^. *GNAS* knockout are also associated with edema, obesity, insulin resistance, and altered energy metabolism, which indicates that this may be a highly pleiotropic genetic target for multiple interrelated physiologies essential to hibernation^76,78^. ()

***ADCY3-*** and ***ADCY7-*** were found to be evolving under relaxed selection in hibernating mammals and are both members of the adenylate cyclase (*ADCY*) family, which play a key role in converting ATP to cyclic AMP (cAMP)^79^. Adenylate cyclase activity has been shown to be suppressed during hibernation (a likely mechanism for energy conservation during hibernation), and relaxed selection on *ADCY7* may indicate that suppressed function of this gene (and potentially other adenylate cyclases) may be a means of bypassing complex regulatory pathways to manage thermogenic responses during hibernation^79,80^. (). *ADCY3* (which regulates fat breakdown in brown adipose tissue (BAT) for heat generation), along with *GNAS* was shown to be highly enriched in brown adipose tissue in hibernating thirteen-lined ground squirrels (*Ictidomys tridecemlineatus*) suggesting it plays a key role in regulating energy expenditure and thermogenesis during hibernation. ***NECAB1^*** encodes a protein involved in calcium signaling in the nervous system and though the mechanism is unknown, it has been identified as being under selection for cold tolerance in chickens^25,81,82^.

***Immune System (2)***

Hibernation significantly suppresses immune function in mammals which increases infection risk^83–86^. While intermittent arousal phases that temporarily restore immune activity are thought to mitigate this decrease, little is known about the molecular mechanisms that may influence this shift in immunity^84^. In these data we recover two genes involved in innate immunity ***NCF4+*** and ***IGSF22⌄*** (evolving slower in hibernators). *NCF4* plays a crucial role in the immune system by generating (ROS) during the respiratory burst in phagocytic cells like neutrophils and macrophages, and dysfunction in this gene can weaken response to pathogens^87,88^. *NCF4* is downregulated in heat stressed birds and hibernating alligators, suggesting that temperature stress may be inhibiting the complement system^89,90^.

**Genes Driven by Bat Evolution**

At least three significant genes recovered in our rate analysis rank very high in the distribution gene recovered when we repeat the same analysis designating bats as the foreground taxa (1st, 3rd, and 6th), and therefore seem to be driven primarily by bat evolution (Table 1). While these are unlikely to be important candidates for hibernation-related adaptive function, they appear to be strongly linked to vision loss (which is well-known in several bat lineages) as two of these genes are associated with eye evolution and the loss/mutation of these genes has been shown cause congenital cataracts (***GJA8^***), and autosomal recessive achromatopsia (***CNGB3^***) ^91,92^. The third gene (***NDST3^***) codes for a heparin sulfate modifying enzyme which has been shown to cause subtle changes in hemostasis but is largely compensated for by paralogs^93^.
