## Supplementary for "Genes Underlying Adaptive Physiological Shifts Among Hibernating Mammals": Supplementary Tables 4-5.docx

**Supplementary Table 4.** Genes showing significant evidence of positive selection (p < 0.05) after correction for multiple tests in BUSTED. Results for several nested tests of selection using BUSTED and RELAX in the HyPhy suite. Foreground species include all ancestral nodes assigned to the foreground via parsimony. Red boxes indicate results which are evidence for signal of selection being exclusive to hibernating lineages, pink boxes indicate results which cannot rule out alternative hypotheses (pervasive selection or selection coming primarily from bat lineages.

| **Gene Name** | **Foreground** | **Test** | **Significant in HF Test** | **Significant in Drop Test** | **P-value w/o Bats** |
| --- | --- | --- | --- | --- | --- |
| ALDH2 | All Hibernators+ Bears Only | BUSTED | no | no | 4.47E-12 |
| RPS6KB1 | ALL Hibernators, no bats | BUSTED | no | no | 1.99E-02 |
| GNAS | ALL Hibernators, no bats | BUSTED | no | no | 4.88E-02 |
| NCF4 | All Hibernators | BUSTED | yes | no | 3.50E-10 |
| PTPN11 | All Hibernators | BUSTED | yes | no | 1.00E-07 |
| NDUFS3 | All Hibernators | BUSTED | yes | no | 1.77E-06 |
| NDUFA12 | All Hibernators | BUSTED | yes | no | 8.88E-05 |
| PPP1CB | All Hibernators | BUSTED | yes | no | 4.67E-03 |
| PPP1R3B | ALL Hibernators, no bats | BUSTED | yes | no | 1.51E-02 |
| AGTR1 | All Hibernators | BUSTED | no | no | 0.120652663 |
| SCT | All Hibernators | BUSTED | no | no | 0.131073405 |
| GCG | All Hibernators | BUSTED | no | no | 0.168901079 |
| NDUFA3 | All Hihernators | RELAX | no | no | 0.5 |
| NDUFB10 | All Hibernators | BUSTED | no | p = 0.05 | 0.445207874 |
| NDUFS5 | All Hibernators | BUSTED | no | p < 0.001 | 0.5 |
| NDUFA2 | All Hibernators | BUSTED, RELAX | no | yes | 0.5 |
| PRKCB | All Hibernators+ Bears Only | BUSTED, RELAX | no | p < 0.001 | 0.5 |
| LRRN1 | All Hibernators | BUSTED | yes | no | 0.5 |

**Supplementary Table 5.** Genes under significant relaxed selection (p < 0.05), after correction for multiple tests in RELAX. Hypothesis free testing is reported for BUSTED, as an equivalent test is not available for RELAX. Foreground species include all ancestral nodes assigned to the foreground via parsimony. Red boxes indicate results which are evidence for signal of relaxed selection being exclusive to hibernating lineages, pink boxes indicate results which cannot rule out alternative hypotheses (pervasive selection or selection coming primarily from bat lineages.

| **Gene Name** | **Foreground** | **Test** | **Significant in HF BUSTED Test** | **P-value w/o Bats** |
| --- | --- | --- | --- | --- |
| NDUFA1 | All Hibernators | RELAX | no | <0.05* |
| NDUFA10 | All Hibernators+ Bears Only | RELAX | yes | <0.05* |
| NDST3* | All Hibernators+ Bears Only | RELAX | yes | <0.05* |
| ADCY7 | All Hibernators+ Bears Only | RELAX | yes | <0.05* |
| ACAT2 | All Hibernators+ Bears Only | RELAX | yes | <0.05* |
| ABCC3 | All Hibernators+ Bears Only | RELAX | yes | <0.05* |
| ACADL | Bears Only, no bats | RELAX | yes | <0.05* |
| ADCY3 | Bears Only, no bats | RELAX | yes | <0.05* |
| ACTB | Bears Only, no bats | RELAX | yes | <0.05* |
| CAMK2A | Bears Only, no bats | RELAX | yes | <0.05* |
| CAMK2G | Bears Only, no bats | RELAX | yes | <0.05* |
| FGFR1 | Bears Only, no bats | RELAX | yes | <0.05* |
| NR1H3 | Bears Only, no bats | RELAX | yes | <0.05* |
| RPS6KA3 | Bears Only, no bats | RELAX | yes | <0.05* |
| RPTOR | Bears Only, no bats | RELAX | yes | <0.05* |
| SLC2A4 | Bears Only, no bats | RELAX | yes | <0.05* |
| SLC4A2 | Bears Only, no bats | RELAX | yes | <0.05* |
| SLC4A4 | Bears Only, no bats | RELAX | no | <0.05* |
| SLC4A5 | Bears Only, no bats | RELAX | yes | <0.05* |
| SMYD2 | Bears Only, no bats | RELAX | yes | <0.05* |
